# Intraocular injection of ES cell-derived neural progenitors improve visual function in retinal ganglion cell-depleted mouse models

**DOI:** 10.1101/114678

**Authors:** Mundackal Sivaraman Divya, Vazhanthodi Abdul Rasheed, Tiffany Schmidt, Soundararajan Lalitha, Samer Hattar, Jackson James

**Author notes:** **Corresponding author:** Jackson James, Ph.D., Neuro Stem Cell Biology Laboratory, Neurobiology Division, Rajiv Gandhi Centre for Biotechnology, Thycaud PO, Thiruvananthapuram, Kerala-695014, India.

## Abstract

Retinal ganglion cells (RGC) transplantation is a promising strategy to restore visual function resulting from irreversible RGC degeneration occurring in glaucoma or inherited optic neuropathies. We previously demonstrated FGF2 induced differentiation of mouse embryonic stem cells (ESC) to RGC lineage, capable of retinal Ganglion Cell Layer (GCL) integration upon transplantation in mice. Here, we evaluated possible improvement of visual function by transplantation of ES cell derived neural progenitors in RGC depleted glaucoma mice models. ESC derived neural progenitors (ES-NP) were transplanted into NMDA (N-Methyl-D-Aspartate) injected, RGC-ablated mouse models and a pre-clinical glaucoma mouse model (DBA/2J) having sustained higher intra ocular pressure (IOP). Visual acuity and functional integration was evaluated by behavioural experiments and immunohistochemistry, respectively. GFP-expressing ES-NPs transplanted in NMDA-injected RGC-depleted mice differentiated into RGC lineage and possibly integrating into GCL. An improvement in visual acuity was observed after two months of transplantation, when compared to the pre-transplantation values. Expression of c-Fos in the transplanted cells, upon light induction, further suggests functional integration into the host retinal circuitry. However, the transplanted cells did not send axonal projections into optic nerve. Transplantation experiments in DBA/2J mouse showed no significant improvement in visual functions, possibly due to both host and transplanted retinal cell death which could be due to an inherent high IOP. We showed that, transplantation of ES-NPs into the retina of RGC-ablated mouse models could survive, differentiate to RGC lineage, and possibly integrate into GCL to improve visual function. However, for the survival of transplanted cells in glaucoma, strategies to control the IOP are warranted.

## Introduction

Glaucoma is a chronic neurodegenerative disease, characterized by progressive, permanent visual loss resulting from the degeneration of retinal ganglion cells (RGCs) (Quigley and Broman, 2006). An increased intra ocular pressure (IOP) resulting from the disrupted dynamics of aqueous humor flow is mainly attributed to the progressive damage of RGC and their axons (Sherwood et al., 2009). Current treatment strategies focus mainly on pharmacological and surgical interventions to control the elevated IOP, which is the prime risk factor for glaucoma (Lee and Higginbotham, 2005). However, the irreversible vision loss as well as RGC degeneration often persists, even when IOP is controlled.

Thus selective replacement of degenerated RGCs to restore lost visual function forms an alternative strategy to regenerate the RGC layer (Chamling et al., 2016). However, vision restoration at translational level in clinics has been hindered by several challenges; such as source of the stem cells, differentiation into appropriate cell type, transplantation approaches, survival and functional integration of cells into the target retina (Hertz and Goldberg, 2013). Additionally, to establish functional neuronal connectivity, optimal microenvironment mediated by the trophic factors is crucial (Johnson and Martin, 2012). Moreover, survival of the transplanted cells also depends on the glaucomatous retinal microenvironment. Although several studies in animal models and isolated clinical trials for cell transplantations have been reported in ocular neurodegenerative diseases (Chamling et al., 2016; Klassen, 2016; Shirai et al., 2016) (Schwartz et al., 2012), results were not substantial to be translated as a therapeutic measure in clinics, owing to the safety concerns, elusive molecular mechanism in establishing the functional connectivity and survival of transplanted neurons (Klassen, 2016). Despite these advances less progress has been made in cell replacement strategies pertaining to RGC loss.

We have previously demonstrated the potential of embryonic stem (ES) cells to differentiate along RGC lineage induced by growth factors, which were capable of integrating into the Ganglion Cell Layer (GCL) of the retina upon transplantation (Jagatha et al., 2009). In the current study, we evaluated restoration of visual function by transplantation of ES cell derived neural progenitors in (i) chemically-induced glaucoma model generated by intravitreal injection of NMDA in C57BL/6 mice and (ii) a pre-clinical model for glaucoma (DBA/2J mice), characterized by high IOP. From the visual behavioural and functional connectivity experiments in NMDA injected RGC-depleted mouse models; we showed that transplanted cells could survive, differentiate to RGC lineage, and possibly integrate into the

GCL resulting in improved visual function. However, the DBA/2J animals did not show any significant improvement in visual acuity behaviour, owing to the degeneration of transplanted cells, which could be resulting from inherent high IOP in the genetic model.

## Materials and Methods Animals

### Animals

All animal experiments were conducted in accordance with the ARVO Statement for the Use of Animals in Ophthalmic and Vision Research. The animal experiments were approved and carried out according to the Johns Hopkins University, Animal Care and Use Committee (No: MO16A212), NIH guidelines, and Institutional Animal Ethics Committee (No: IAEC/215/JAC/2013) of Rajiv Gandhi Center for Biotechnology. All mice were housed in standard animal facility cages with access to food and water *ad libitum* with 12h light/dark cycle.

### Generation of NMDA injected glaucoma model

C57BL/6 mice strain aged 6-8 weeks was used for generating RGC ablated model by injecting NMDA intravitreally. Animals (n = 20) were anesthetized with 0.3 - 0.5ml of Avertin (20mg/mL) and 2μl of 100μM NMDA was injected using graded glass micropipettes with a fine tip aperture into both eyes. The mice were analyzed for visual function behavioral tests (n =10), and further subjected to immunohistochemical analysis for RGC loss (n = 5), and anterograde tracing of RGC axonal projections (n =5) after 1 week, in comparison to the non-injected controls (n =10). For transplantation experiments remaining NMDA injected animals (n=10) were used.

### DBA/2J glaucoma model

The DBA/2J mice, pre-clinical model of glaucoma (8 months old, n = 6) exhibiting elevated IOP resulting from point mutations in two genes, *Gpnmb* and *Tyrp1* (Wang et al., 2000; de Melo et al., 2003) were also used for transplantation studies. DBA/2J-*Gpnmb^+^* genetic control mice strain (8 months old, n =3) with normal IOP was used as corresponding experimental control (Both mouse strains were a kind gift from Nick Marsh Armstrong’s Laboratory, Johns Hopkins University, School of Medicine, Baltimore, MD).

### Transplantation

We used stable GFP expressing CE3 ES cells, (ATCC SCRC-1039) for our differentiation and transplantation experiments. The transplantation experiments were performed as per our previous report (Jagatha et al., 2009). Briefly embryoid body (EB) induced from CE3 ES cells were partially differentiated on poly-D-Lysine (150 μg/ml) and Laminin (1 μg/ml) substrate for 2 days in neuron differentiation medium [DMEM/F12 supplemented with 1% N2 supplement (Invitrogen), 0.5% FBS, Heparin (2μg/ml) and FGF2 (10 ng/ml) (Chemicon)]. The partially differentiated EBs were trypsinized and plated on to uncoated plates in medium consisting of DMEM/F12 supplemented with 1% N2 supplement, Heparin (2 μg/ml) and FGF2 (10 ng/ml) for 6 days to generate ES cell derived neural progenitor (ES-NP) cells. These ES-NPs were dissociated into single cell suspension (1×10^6^ cells/μl in 1X PBS and 10 ng/ml FGF2) and were used for transplantation experiments. Approximately ~1million (ES-NP) cells were transplanted intravitreally in NMDA injected-RGC depleted mice models (n = 10) and DBA/2J mice, pre-clinical glaucoma mice model (n = 6). Briefly, a pulled capillary glass micropipette connected to a micro-injector was injected into the eye near the equator and retina to deliver 1-2μl cell suspension (~1×10^6^ cells) into the vitreous. The behavior experiments for visual function were tested after two months of transplantation and were compared with prior to that of transplantation. Further the transplanted animals were sacrificed for immunohistochemical characterization of retina.

### Behavioral Analysis

#### Optokinetic tracking experiments

We used an optokinetic tracking system (OptoMotry, Cerebral Mechanics Inc.) to evaluate the visual acuity by measuring the image-tracking reflex of mice (Prusky et al., 2004). Visual stimuli are projected on to the monitors so that a virtual cylinder with rotating gratings (sine wave grating) is produced. The mouse was allowed to track the grating and the tracking response was recorded. All acuity thresholds were determined using the staircase method with 100% contrast (Güler et al., 2008).

#### Light Avoidance Experiment

The innate aversion of mice to the brightly illuminated area was utilized here to evaluate the visual responses. The light avoidance test was conducted using a light-dark box consisting two equal sized compartments (20cm × 20cm). The light compartment was kept eliminated at a luminosity of 600 lux. An opening (7.5 × 7.5 cm) was located in the wall between the two chambers allowing free access between the light and dark compartments. The mice was placed in the light chamber to explore the box for five minutes and their movement across the chambers was tracked by computerized video tracking system (Anymaze, Stoelting, USA). The fraction of time spent by the mice in the open illuminated space is noted for calculating the percentage of time spent in light zone.

### Anterograde tracing of RGC axonal projections in NMDA model

NMDA injected mice (n = 5) and non-injected controls (n = 5) were anesthetized as mentioned previously and 2μl of cholera toxin B subunit conjugated with Alexa Fluor 488 (CTB-488, Invitrogen, CA) was injected intravitreally to track the RGC axonal projections. Three to four days after injection, the retina and brain were fixed by perfusing with 4% Paraformaldehyde (PFA). The brain samples were dissected and coronal sections were taken for further analysis. RGC axonal targets in the brain such as Suprachiasmatic Nuclei (SCN), Lateral Geniculate Nuclei (LGN) and Superior Colliculus (SC) were analyzed for RGC axonal projection innervations. The levels of fluorescence of maximum RGC projections in brain regions and total fluorescence levels were measured with Image-J software (http://rsbweb.nih.gov/ij/) as described previously (Karali et al., 2007).

### Immunohistochemistry

For immunohistochemistry, whole eye ball has been fixed overnight in 4% paraformaldehyde, and the fixed tissues were dehydrated in 30% sucrose, embedded in OCT (Optimal Cutting Temperature; Tissue-Tek), and 12-μm cryostat sections were cut through the retina. Immunofluorescence analyses were carried out on tissue sections where, the sections were blocked in 1X PBS containing 5% NGS and 0.2% Triton X-100 followed by an overnight incubation in primary antibody at 4˚C, washed in 1X PBS and incubated with appropriate secondary antibody for 45 min, and mounted with a mounting medium (Fluoromount-G, Electron Microscopy Science, USA) and counterstained with DAPI for visualizing the nucleus. The primary antibody used was mouse anti-GFAP (1:200, Sigma), and secondary antibody was conjugated to Cy3 (1: 400 dilution, Jackson immuno-research).

### Whole mount Retina Immunostaining

To evaluate the RGC loss, NMDA injected animals (n = 5) and non-injected control animals (n = 5) were sacrificed and the eyes were fixed in 4% PFA for 30 minutes followed by dissection of retina. The retina was again fixed in 4% PFA for one hour followed by blocking in 0.6% blocking solution. The whole mount retina was incubated with rabbit anti-Brn3a primary antibody (1:500, Chemicon) at 4°C for 2-3 days followed by washing with 1X PBS three times for 30 minutes each. Goat anti-Rabbit IgG (H+L), Alexa Fluor 594 (1:400, Molecular Probes) was used as secondary antibody and incubated for two days at 4°C.

In order to assess functional integration of transplanted cells, we analyzed the expression of an immediate early gene c-Fos in transplanted NMDA injected mice housed in normal 12:12 light-dark cycle. Mice were given a light pulse of 500 lux intensity for 30 minutes after which mice were placed back in dark for 90 minutes. Animals were subsequently sacrificed, enucleated and whole mount retinal preparation were made as described earlier. The whole mount retinas were then incubated with rabbit anti-c-Fos antibody (1:10,000, Calbiochem) at 4°C for 2 days followed by secondary staining with Goat anti-Rabbit IgG (H+L), Alexa Fluor 594 (1:400, Molecular Probes). To confirm the RGC lineage of transplanted cells, immunostaining with the RGC marker, Calretinin (García-Crespo and Vecino, 2004; Lee et al., 2010) (Rabbit anti-Calretinin, 1:500, Chemicon) was done. The sections were mounted on microscopic slides in mounting medium (Vectashield, Burlingame, CA) with DAPI for nuclear staining. The slides were viewed and imaged in an upright fluorescent microscope (Carl Zeiss, Oberkochen, Germany).

### Wheel Running Activity

Both control and NMDA injected mice were placed in cages with a 4.5-inch running wheel, and their activity was monitored with VitalView software (Mini Mitter, OR). Photoentrainment experiments were done under a 12 hours light: 12 hours dark cycle. Initial light intensity was provided by Philips Daylight deluxe fluorescent lamps. The periods in constant darkness and constant light were calculated with ClockLab (Actimetrics, IL).

### Statistical Analysis

All data were obtained by at least three independent experiments and represented as Mean ± SD. Statistical significance between two groups was calculated by unpaired *t-tests* assuming two-tailed distribution and equal variances. *P* value <0.05 was considered as statistically significant.

## Results

### Visual acuity of the NMDA injected mice is significantly reduced along with RGC loss

The RGC ablated mice model was generated by intravitreal injection of NMDA (100μM) to induce excitotoxic damage to RGCs. After one week of NMDA injection, there was ~50% reduction in the number of Brn3a positive RGCs in the whole mount retina (33.85 ± 4.71) when compared to control animals with no NMDA injection (57.74 ± 4.66, p<0.001, Fig.1 A-F&G). In order to understand the possibility of whether the NMDA injected retina could regenerate the lost RGC’s induced due to excitotoxicity, we also analysed the retina after 40 days of NMDA injection and observed that the loss is irreversible (**Fig. S1A-C**). Visual acuity of the NMDA injected mice was tested by its ability to track black and white gratings on a virtual rotating cylinder using an optokinetic tracking system (Fig.1 H). Our results showed that there was a significant reduction in visual acuity (cycles/degree) of NMDA injected animals (0.209 ± 0.053, p < 0.001, control; n = 10, before and after NMDA injection; n = 10, (Fig.1 J), compared to the pre-injection controls 0.427 ± 0.08. A light avoidance test (Fig.1 I) also supported the above observation, where the pre-injection controls avoided bright light and spent only ~17% of the time in light zone. The NMDA injected animals showed comparatively less light avoidance behaviour and spent ~43%, (p < 0.001) of time in the light zone, (Fig.1 K). Similar results were also observed after 40 days of NMDA injection indicating that the animals have lost visual functions permanently with no recovery (**Fig. S1D**). These results clearly suggested that ablation of RGCs by NMDA injection, leads to impairment in visual functions.

**Figure 1:**
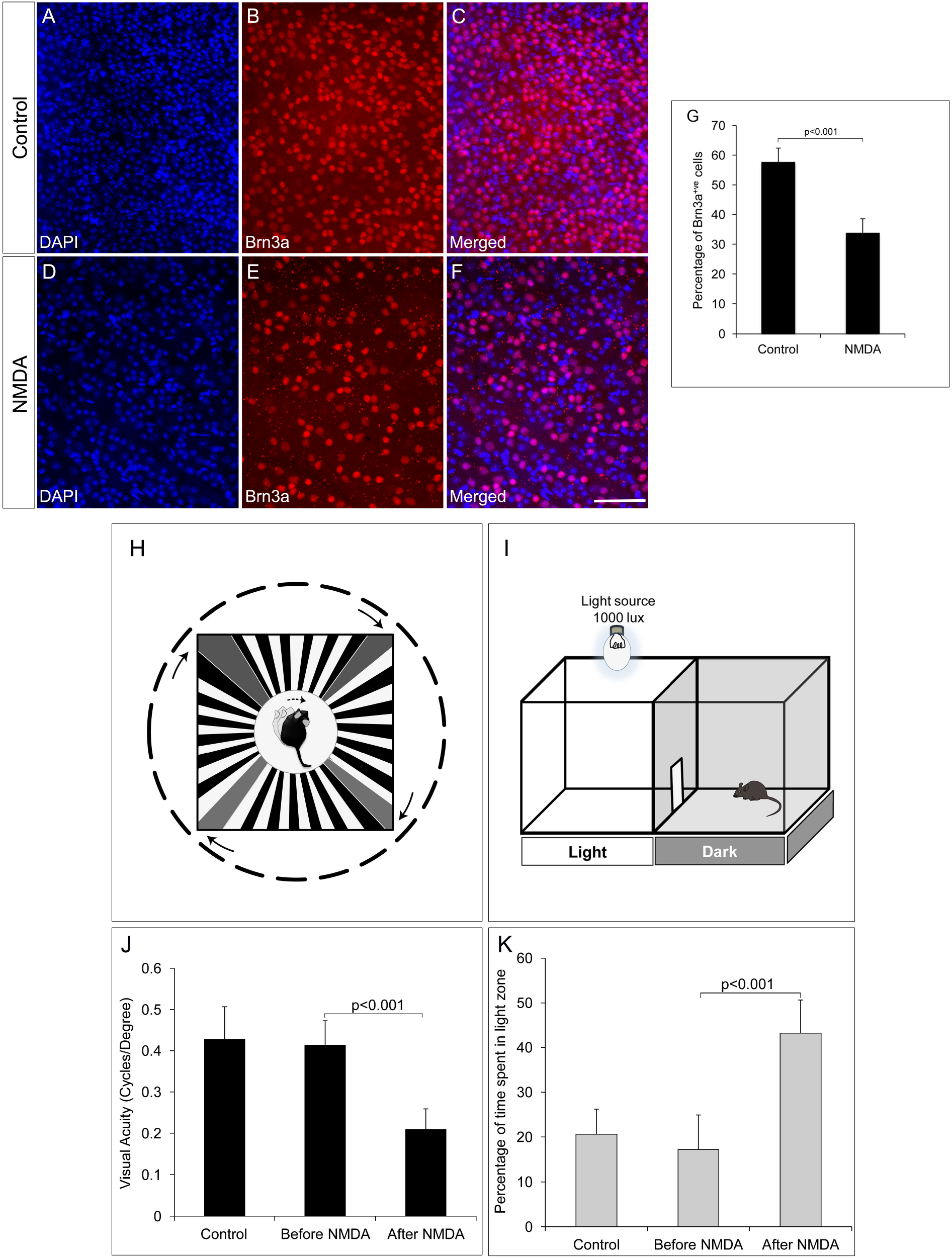
NMDA-injected animals show reduction in number of RGCs and consequent visual function deficits. A-F) Brn3a whole mount retina immunostaining showing reduced number of Brn3a positive cells in NMDA-injected animals compared to control animals. G) Graph represents quantitative analysis showing the percentage of Brn3a positive cells. H-I) Schematic of Optomotry and light avoidance test set up. J) Visual acuity of NMDA-injected animals is significantly low compared to that of control animals. K) NMDA-injected animals also showed less light avoidance behavior compared to that of control animals. Data are expressed as Mean ± SD. Number of animals used for behavioral experiments; Control = 10 and NMDA injected = 10. Scale = 100μm.

We further investigated RGC innervation of brain targets, both in image forming visual function centres (LGN and SC) and non-image forming visual function centres (SCN). Anterograde tracing experiments with intravitreal injection of Cholera toxin B-488 was performed to track the RGC axonal projections. All images were taken with the same exposure time. RGC innervations into the dorsal LGN of NMDA injected animals (218.00 ± 7.86) was significantly reduced compared to that of control animals (682.00 ± 42.41, p < 0.001, NMDA injected mice; n = 5, non-injected controls; n = 5, Fig.2 A-D&E). Similarly, SC also had significantly less RGC innervations in NMDA injected animals (274.24 ± 97.32) compared to that of controls (2502.85 ± 6.77, p < 0.001, Fig.2 F-I&J). However, the RGC innervations to the SCN, the brain region responsible for regulating the circadian rhythm were intact (Control = 634.47 ± 53.53, NMDA=404.28 ± 80.65, p < 0.21, NMDA injected mice; n = 5, non-injected controls; n = 5, Fig.2 K-N&O). Although the RGC innervations were intact, there was an observed decrease in innervations which was not statistically significant (Fig. 2O). These observations were in agreement with our results where the NMDA injected animals showed drastically reduced visual acuity which might be due to the lesser RGC innervation in the image forming centres of the brain as of the RGC loss. The intact innervation in the SCN was in agreement with the reports that the NMDA injection does not affect ipRGCs (DeParis et al., 2012). The circadian rhythm pattern, as evidenced from the wheel running activity further confirmed no differences between the NMDA injected mice and controls (**Fig. S2**). We thus used animals, after 7 days of NMDA injection, as RGC ablated model for the transplantation studies.

**Figure 2:**
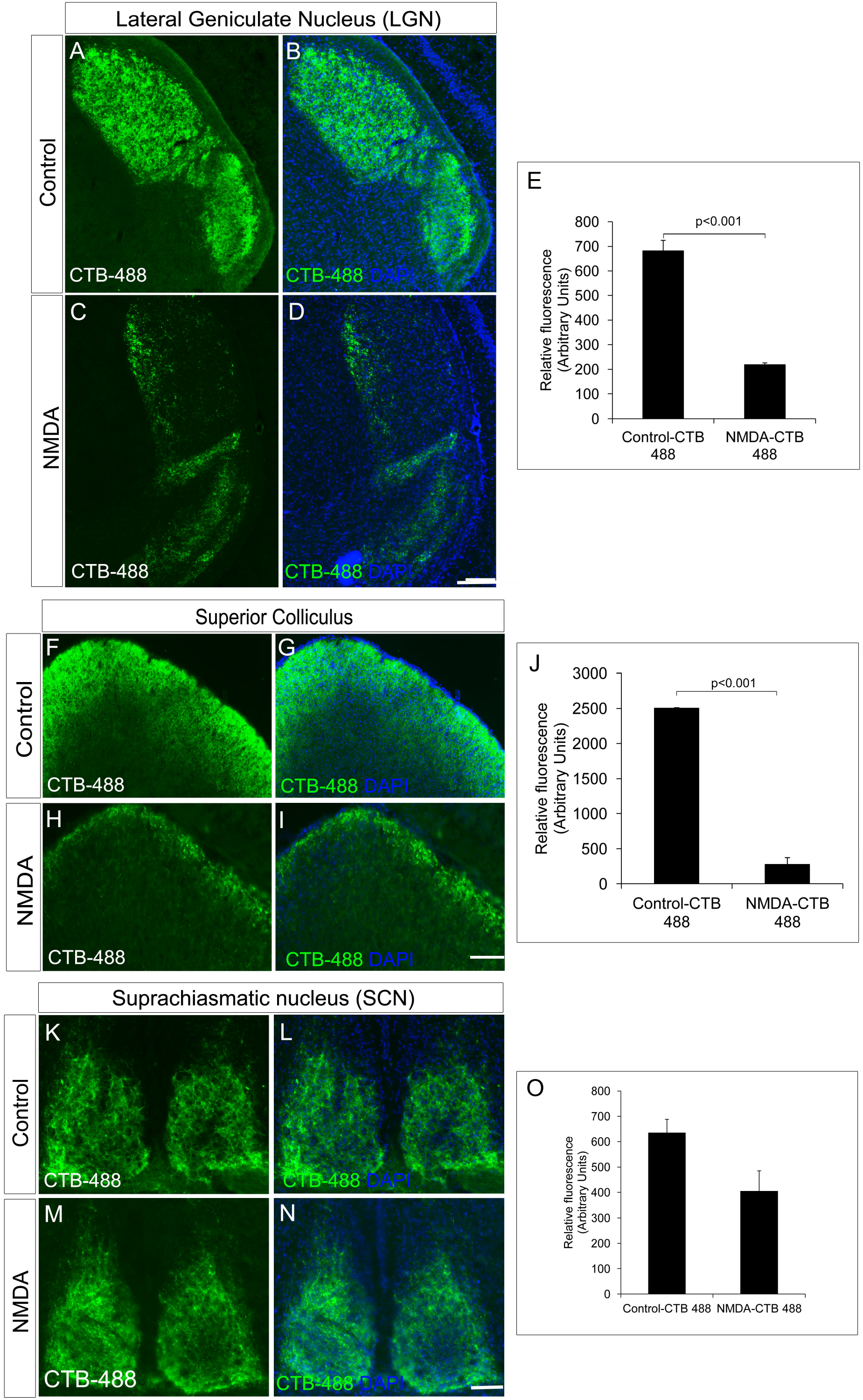
NMDA-injected animals showed a significant reduction in RGC projections into the image forming centres in the brain, LGN and SC but not in non-image forming centre such as SCN. A-D&E) CTB-488 shows that axonal projections into the dorsal LGN are significantly reduced in the NMDA-injected animals compared to that of the control animals. F-I&J) Anterograde tracing experiment showed significant reduction in RGC projections to Superior Colliculus in NMDA-injected animals. CTB-488 staining, tracking RGC projections into SC was significantly low in NMDA-injected animals compared to that of control animals. RGC projections into non-image forming brain centres are not affected in NMDA-injected animals. K-N&O) RGC projections into SCN of NMDA-injected animals are almost intact similar to that of control. All images were taken with the same exposure time. Data are expressed as Mean ± SD. Scale = 100μm.

### ES-NPs transplantation in RGC depleted mice improved visual function

We have previously reported that FGF2-induced ES-NPs were able to differentiate along RGC lineage evidenced by the expression of RGC regulators and markers (Jagatha et al., 2009) (**Fig. S3 A-N**). Further we have also shown that ES-NPs, after exposure to FGF2, were capable of integrating and differentiating into RGCs *in vivo* upon transplantation (Jagatha et al., 2009). In the present study we adopted the same strategy. The FGF2 treated, differentiating ES-NPs (GFP expressing) were transplanted intravitreally into the NMDA injected-RGC depleted mice model. The animals were subjected to behavioral tests once again, prior to transplantation and then two months after. Significant improvement in visual acuity was observed in transplanted animals as evidenced from optokinetic response, compared to NMDA injected animals, before transplantation (Transplanted = 0.233 ± 0.07, NMDA injected, prior to transplantation = 0.138 ± 0.07, p < 0.05 Fig.3 A). Though, we observed an improved visual function in NMDA model, we could not observe integration of transplanted cells into the optic nerve.

**Figure 3:**
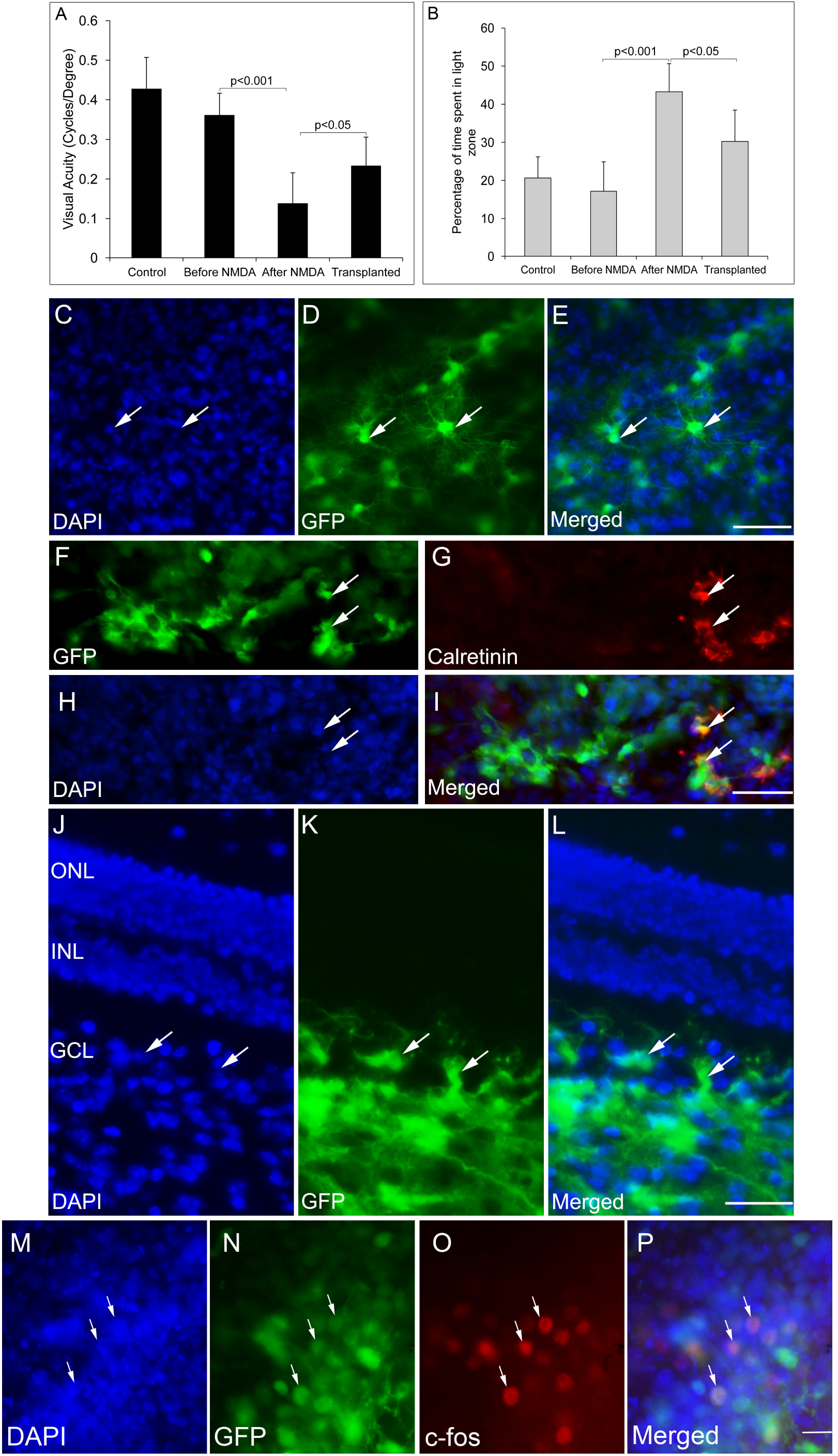
GFP-expressing ES-NPs showed extensive integration and functional connectivity in transplanted NMDA-injected mouse models. A) Transplanted, NMDA-injected animals showed improvement in visual functions. Visual acuity of transplanted, NMDA-injected animals is significantly increased compared to that of NMDA-injected animals prior to transplantation. B) Transplanted, NMDA-injected animals also showed improved light avoidance behavior compared to that of NMDA-injected animals prior to transplantation. C-E) Whole mount immunostaining of retina showed extensive integration of GFP-expressing cells with typical neuronal dendritic processes indicating RGC morphology. F-I) Transplanted GFP-expressing cells co-expressed RGC marker Calretinin. J-L) GFP-expressing ES-NPs integrated into the GCL of NMDA-injected animals as shown by the immunostaining of retinal sections. M-P) GFP-expressing cells integrated into the retina after transplantation expressed c-Fos, an immediate early gene upon light induction confirming functional integration with photoreceptor expressing cells. Data are expressed as Mean ± SD. Number of animals used for behavioral experiments; Control = 10, NMDA-injected (before transplantation) and Transplanted = 10. Scale = 100μm.

In the light avoidance test, animals after transplantation spent less time (90.7 ± 24.8 s, 30%) in the light zone compared to those prior to transplantation (129.7 ± 22.2 s, 43%), indicating a significant improvement (p < 0.05) in light responsiveness. However, the noninjected control animals spent only 17% (51.5 ± 23.2 s) of the time in light zone, exhibiting highest light avoidance behavior (Fig.3 B). Thus the results indicate ES-NPs transplantation has shown improvement in visual behavioral tests suggestive of partial restoration of vision.

To confirm the integration of the transplanted cells into the GCL of host retina, the animals were sacrificed after behavioral analysis and whole mount retinal staining was done. Extensive integration of GFP expressing cells throughout the surface of retina with distinct RGC morphology was observed (Fig.3 C-E). The RGC lineage was also confirmed by immunostaining with the RGC marker, Calretinin (Fig.3 F-I). Retinal sections also showed integration of GFP expressing cells in the RGC specific GCL (Fig.3 J-L).

To test whether the transplanted cells form functional neural circuits with the photoreceptor expressing cells, expression of c-Fos, an immediate early gene, was probed in the transplanted cells, in response to light induction. The transplanted GFP-expressing cells showed c-Fos expression, suggestive of getting activated by the light stimulus (Fig.3 M-P). Thus the results are suggestive of transplanted cells capable of being functionally integrated into the host retinal circuitry by forming connections with the photoreceptor expressing host cells. Although the number of cells co-expressing c-Fos was less in number, this was a very promising result, which substantiates our results of improved visual acuity evidenced from behavioural experiments (Fig.3 A, B).

### ES-NPs transplantation in DBA/2J mice did not show significant improvement in visual functions

We further extended our transplantation protocol to the pre-clinical glaucoma model, DBA/2J mice of 8 months of age. The DBA/2J mice had deficits in visual functions and could hardly track the gratings in optokinetic tracking system when compared to the corresponding DBA/2J-*Gpnmb^+^* genetic control (DBA/2J-*Gpnmb^+^* genetic control = 0.276 ± 0.03, DBA/2J = 0.137 ± 0.02, p < 0.001 Fig.4 A). Further DBA/2J mice also showed less light avoidance behaviour compared to the DBA/2J -*Gpnmb^+^* genetic control animals confirming their visual deficits (Control 45.57 ± 11.64 s, 15%, DBA/2J 174.08 ± 23.32 s, 58 %, p < 0.001, Fig.4 B).

**Figure 4:**
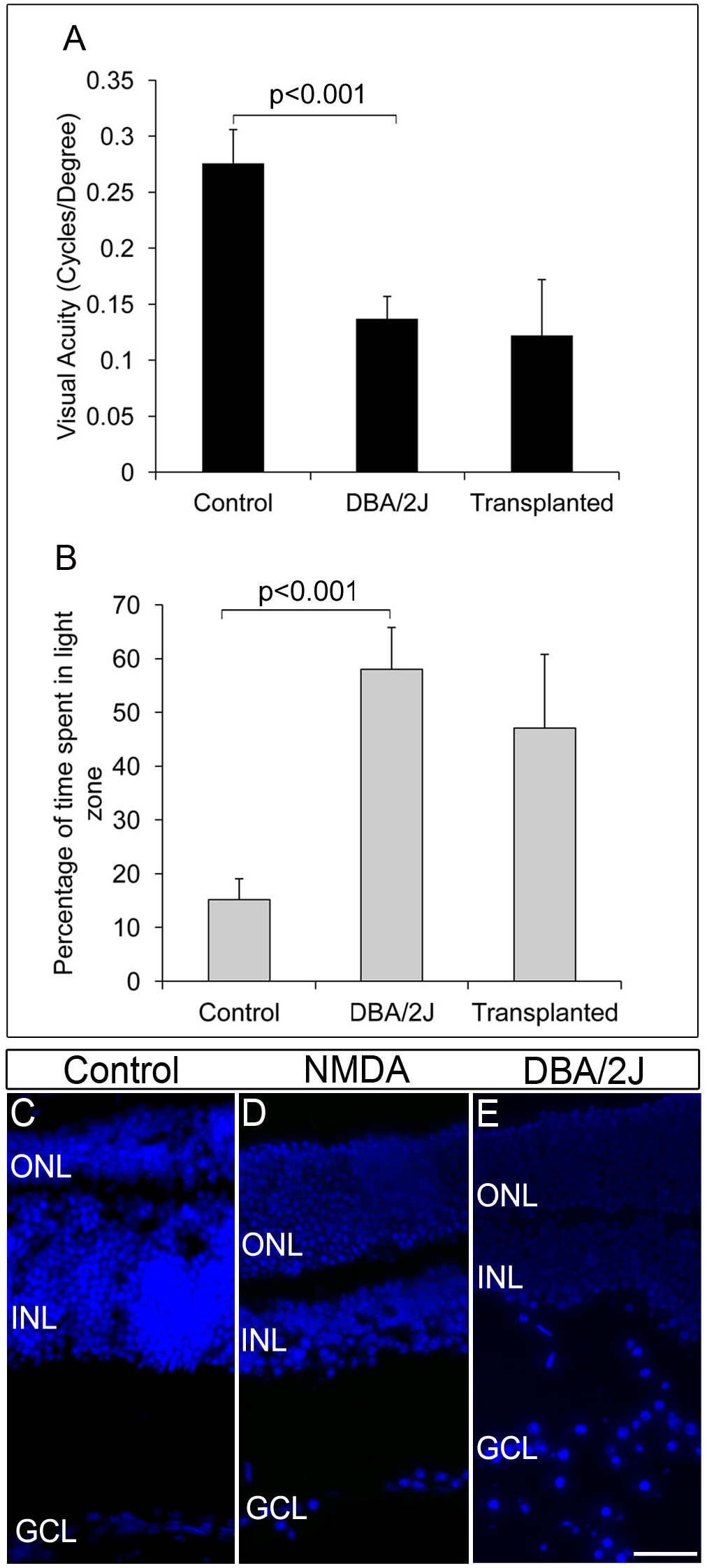
DBA/2J animals did not show any improvement in visual functions after transplantation. A) Though there was a significant decrease in visual acuity of DBA/2J animals compared to that of control (p<0.001), no significant improvement in visual acuity was seen after transplantation in DBA/2J animals. B) Light avoidance behavior was also very poor in transplanted animals indicating that there was no significant vision recovery after transplantation in DBA/2J animals. Data are expressed as Mean ± SD. Number of animals used; Control = 3, DBA/2J (before transplantation) and Transplanted = 6. C) DAPI staining of control eye section showing intact GCL layer of the retina D) NMDA-injected animals showed disorganized GCL layer with less number of RGCs compared to that of control wild type animals. E) Pre-clinical glaucoma model, DBA/2J also showed a highly disorganized GCL layer with scattered RGCs. Scale = 100μm.

FGF2-induced GFP-expressing ES-NPs were transplanted into the visually challenged DBA/2J animal. Visual functions by behavioural experiments were tested after two months of transplantation. As evidenced from optokinetic response and light avoidance behaviour, visual acuity in DBA/2J animals did not improve after transplantation (Fig.4 A, B). Moreover DBA/2J mice had GCL with very low number of RGCs as that of NMDA injected mice (Fig.4 D) except which are highly disorganized and dispersed irregularly throughout (Fig.4 E) compared to that of controls (Fig.4 C). We speculate that the elevated IOP inherent to the DBA/2J mice might induce oxidative stress (Moreno et al., 2004) on the transplanted cells, preventing them from proper differentiation and innervation, limiting the improvement in visual acuity. Therefore, by comparing the NMDA and DBA/2J models, it appears that control of IOP is much more crucial for the transplanted cells to integrate efficiently for functional recovery.

## Discussion

Large number of studies has been reported, where photoreceptor transplantation in appropriate disease models showed partial restoration of vision (Yang et al., 2010; Tucker et al., 2011; Pearson et al., 2012; Santos-Ferreira et al., 2015; Chamling et al., 2016). However, data is limiting with RGC transplantation in glaucoma models. Here, we have evaluated the functional visual restoration upon transplantation of ES-NPs in two glaucoma animal models; NMDA-induced RGC-depleted model and DBA/2J mice, a standard pre-clinical genetic glaucoma model (Anderson et al., 2001).

Our transplantation results in the NMDA-induced RGC-depleted mouse models demonstrated possible integration of ES cell-derived RGC like cells into GCL layer with proper RGC morphology and RGC marker expression. Further co-expression of c-Fos with GFP-expressing cells in transplanted animals upon light induction demonstrated an improved functional integration into the GCL. In addition, behaviour experiments to evaluate visual acuity also augmented the functional vision restoration, although partial, in the transplanted animals. Similar observations have been reported previously, which showed an improved retinal scotopic wave function in NMDA-injected rats after transplantation of human Müller glial derived RGCs (Singhal et al., 2012). Improvement of visual functions evidenced by electroretinogram (ERG) and visual evoked potential (VEP) following the transplantation of retinal progenitor cells in rat models of retinal degeneration was also reported (Li et al., 2014; Qu et al., 2015).

Nevertheless, in our study integration of transplanted cells into optic nerve was not observed, which was in line with reports where the transplanted cells were seen to extend processes into the RGC layer, but not towards the optic nerve (Singhal et al., 2012; Hertz et al., 2014). Currently, we do not have a clear cut evidence to prove the observed functional recovery, even though we do not see the extension of axons from the transplanted cells to the brain visual function. We speculate that the transplanted cells could possibly integrate into the GCL layer and may establish local interneuron connectivity and also secrete neurotrophic factors exerting a protective effect on optic nerve, preventing further neuronal loss and improve visual functions (Singhal et al., 2012; Becker et al., 2016). Moreover, NMDA-induced model showed only ~50% RGC depletion suggesting a possible contribution of the transplanted cells in the survival of resident RGCs in the retina and their axons providing a repairing effect on optic nerve that innervate the visual centres in the brain. However, the present study was limited by the neuronal tracking to show the restored neuro-anatomy in the visual centres of brain. Further electrophysiological evidences were also lacking to support the recovery, but the improved visual functions evidenced from behavioural studies along with the observations from c-Fos expression results showed possible functional integration of the transplanted cells proving intact functional anatomy. Though this integration was promising the number of c-Fos co-expressing cells were less, which could be attributed to the lesser RGC yield after FGF2 induced differentiation (Jagatha et al., 2009). This also limited us from quantitating the c-Fos expression. Another possibility for the observed reversal in the visual function could be due to cytoplasmic transfer as observed with photoreceptor precursor transplantation resulting in the survival of the native RGCs (Pearson et al., 2012). However this possibility can be ruled out for RGC integration in GCL, as we could observe single nuclei in the integrated cells (Fig.3 C-L). We also do not totally rule out the role of FGF2 in bringing about the observed reversal since in our experiments we used a FGF2 control and not a sham control. Although such a possibility is very remote since the injected FGF2 would be removed from the system within few hours of transplantation. These experiments are planned to completely rule out this possibility.

Notably, a recent report showed integration of transplanted primary mice RGCs in uninjured rat retina, where axons extended to visual centres in brain with electrical excitability and light response (Venugopalan et al., 2016). The uniform population of primary RGCs from developing retina used for the transplantation in a non-pathological model and the factors derived from primary donor RGCs might have contributed for this integration (MacLaren et al., 2006; Hertz et al., 2014). Therefore, to test long range arborisation to visual centres after transplantation, future studies are warranted for precise RGC fate acquisition from pluripotent stem cell source with good yield and using pure population of differentiated RGC for transplantation.

Meanwhile, the DBA/2J animals did not show any significant improvement in vision after transplantation. This could be attributed to the inherent high IOP in DBA/2J animals resulting in the degeneration of the transplanted cells owing to the oxidative stress (Moreno et al., 2004). Since the GCL of retina is highly dispersed in these mice models compared to NMDA GCL which is not disorganised, it can be speculated that the transplanted cells were not able to integrate. Thus the dispersed nature of the GCL may limit the scaffolding support to the transplanted cells. Moreover, the transplantation was done at the age of 8 months, where RGC degeneration occur maximum because of high IOP implying the requirement of pharmacological IOP alleviation for effective transplantation. However, our study was limited by transplantation experiments in DBA/2J controlled for IOP. Therefore, for the survival of transplanted cells in glaucoma, strategies to control the IOP are warranted.

Although, there was a recovery of visual function in the NMDA model, it can be considered as the recovery from the pathological state of glaucoma by replenishment of cells.

Since elevated IOP forms a critical characteristic feature of glaucoma and the NMDA model represents only a static glaucomatous stage of degenerated RGC population. Thus, the ocular disease phenotype modelled in the animal plays a significant role in evaluating the efficacy of the cell transplantation. While we have seen a positive outcome, by transplantation of ES cell-derived RGCs into RGC depleted model, we cannot rule out the role of inflammatory response that could also evoke the secretion of growth factors which could also mimic a similar result (**Fig.S1 E, F**). Previous reports have demonstrated that Interavitreal injection of zymosan (IVZ) causes inflammation which in turn stimulates RGC survival and axon regeneration (Ahmed et al., 2010). Similarly, optic nerve crush injury and lens injury also stimulated axon regeneration (Leon et al., 2000; Lorber et al., 2012). Furthermore, it is known that Müller glia cells can de-differentiate, proliferate, and become neuronal progenitors in acutely damaged retinas, which make it a potential source of progenitor-like cells for retinal regeneration (Fischer and Reh, 2003). The neurogenic potential of injury-activated Müller cells to generate retinal neurons (Das et al., 2006) and partial restoration of RGC function *in vivo,* following its transplantation has been reported (Singhal et al., 2012). Moreover, FGF2 has a prominent role in transdifferentiation of Müller glia cells and proliferation of Müller glia-derived progenitor-like cells (Fischer and Reh, 2003). However, possible contribution of FGF2 in promoting the resident retinal stem cells or excitotoxic injury activated Müller cells to coax towards the RGC lineage warrants further studies. Albeit, we obtained an improvement in visual function after transplantation, the results should be cautiously interpreted, owing to multitude of confounding conditions influencing the functional recovery. Therefore, future studies with more controlled conditions should be implemented to rule out these confounding effects

Altogether, our results showed; the prospect of cell replacement therapy in glaucomatous condition, requirement of appropriate animal models mimicking the pathological human conditions, and advanced transplantation strategies to promote the survival of the transplanted cells. Our findings also point towards the drawbacks that could compromise stem cell therapy where the pathogenic mechanism is still active in the host environment, that is also the case with other ocular degenerative diseases such as age related macular degeneration.

## Authors’ contributions

MSD, JJ and SH conceived and designed the experiments; MSD, performed the experiments; MSD, VAR and TS collected, analyzed and interpreted data. SL generated supplementary figure S1. The manuscript was written through contributions of all the authors. All authors have given approval to the final version of the manuscript.

## Acknowledgements

The authors thank Dr. Ani V Das for her critical reading and advice on this manuscript, Dr. Rejji Kuruvilla, Dr. Haiqing Zhao and the mouse tri-lab members for suggestions and advice for the experiments. The authors also thank, Dr. Nick Marsh Armstrong, School of Medicine, Johns Hopkins University for kindly providing DBA/2J and DBA/2J-*Gpnmb^+^* mice for the experiments.

## List of abbreviations

RGCs: Retinal Ganglion Cells; ES cell: Embryonic Stem Cells; ES-NP: Embryonic stem cell derived Neural Progenitors; IOP: Intra Ocular Pressure; NMDA: N-Methyl-D-Aspartate; GCL: Ganglion Cell Layer; SCN: Suprachiasmatic Nuclei; LGN: Lateral Geniculate Nuclei; SC: Superior Colliculus; ERG: Electro Retinogram; VEP: Visual Evoked Potential

